# Dingent: An Easily Deployable Database Retrieval and Integration Agent framework

**DOI:** 10.64898/2026.03.17.712026

**Authors:** Demian Kong, Shaoqi Bei, Yueyue Wu, Bixia Tang, Wenming Zhao

## Abstract

AI-driven data search and integration represent an emerging research direction. Although several LLM-based backend frameworks and agentic frameworks have emerged, significant gap remains in developing a one-stop, configurable agent framework that supports various data sources and provides a web interface for efficient data retrieval using natural language. To address this, we present Dingent, a novel and configurable agent framework that facilitates data access from various resources and enables the flexible constructions of agent applications. We demonstrate its capabilities across three distinct application scenarios, achieving promising results. The Dingent framework can be readily applied to other fields, such as earth sciences and ecology, to facilitate data discovery.

## Introduction

AI-driven retrieval applications for biological databases represent an emerging research direction. The objective of these application is to enable more efficient use of complex biological data and to provide natural language interfaces for querying. By integrating information from disparate sources, they could generate coherent insights.

Currently, several LLM-based solutions have emerged, which can be broadly classified into three categories. The first includes open-source frameworks such as LangChain (1) and LlamaIndex (2), which are specifically designed for developing LLM-based agents and support various data storage engines, including MySQL and Elastic Search. The second category encompasses intelligent applications such as AI-HOPE (3), MRAgent (4), DeepPGDB (5), and BIODISCOVERYAGENT (6) that demonstrate the potential of AI-driven agents in data retrieval and integration across diverse domains, including disease, multi-omics data integration and experiment design. Among these, the integrated agent platform Biomni (7) provides access to 59 databases and 150 tools across 25 biomedical domains. Finally, agentic frameworks like AutoGen (8) and CrewAI (9) focus on multi agent collaboration and planning, showing strong capabilities in complex reasoning tasks. However, several limitations still exist. First, open-source frameworks like LangChain require programming expertise for development, which limits their accessibility for many life science researchers. Second, most agent-specific retrieval applications (e.g., MRAgent) suffer from a lack of flexibility and generalizability, which hinders their adaptation to evolving research needs. Furthermore, agent frameworks like AutoGen are not designed for building end-to-end, UI-driven applications. There is still a gap in developing a simple, user-friendly, one-stop agent framework that supports various data storage engine configurations and provides a web interface for data retrieval to facilitate data discovery.

To bridge this gap, we present Dingent, a novel and configurable agent framework that offers simplified access to diverse biological data resources. Its flexible architecture allows researchers without programming or AI expertise to integrate through multiple databases through simple configuration. Featuring an intuitive web interface, Dingent enables users to ask questions in natural language. The AI agent interprets each query, dynamically retrieves relevant data, and combines the result into clear insights. Crucially, life scientists can customize the application by adding new data access methods through configuration files.

## Method

### The overview of Dingent

Dingent offers a straightforward and efficient framework for generating agent applications through configuration, which significantly streamlines development. Specifically, an admin configuration page is used to configure the LLM, access data sources, and orchestrate their execution. Once the configuration is complete, a user chat interface for data retrieval is ready to use. The request handling is implemented as FastAPI services, with various services initiated at runtime. The agent execution engine parses user questions, orchestrates data retrieve tasks, calls appropriate plugins to execute them, and summarizes the results. Currently, Dingent facilitates access to various data resources, including MySQL, ChromaDB, Elastic Search, alongside custom data resources. It also supports the construction of various application scenarios, such as single-source, multiple-source, and associated-source retrieval. Dingent is accessible as a Python library, which can be installed from our GitHub repository. The overall architecture is illustrated in Figure 1.

**Figure 1.**
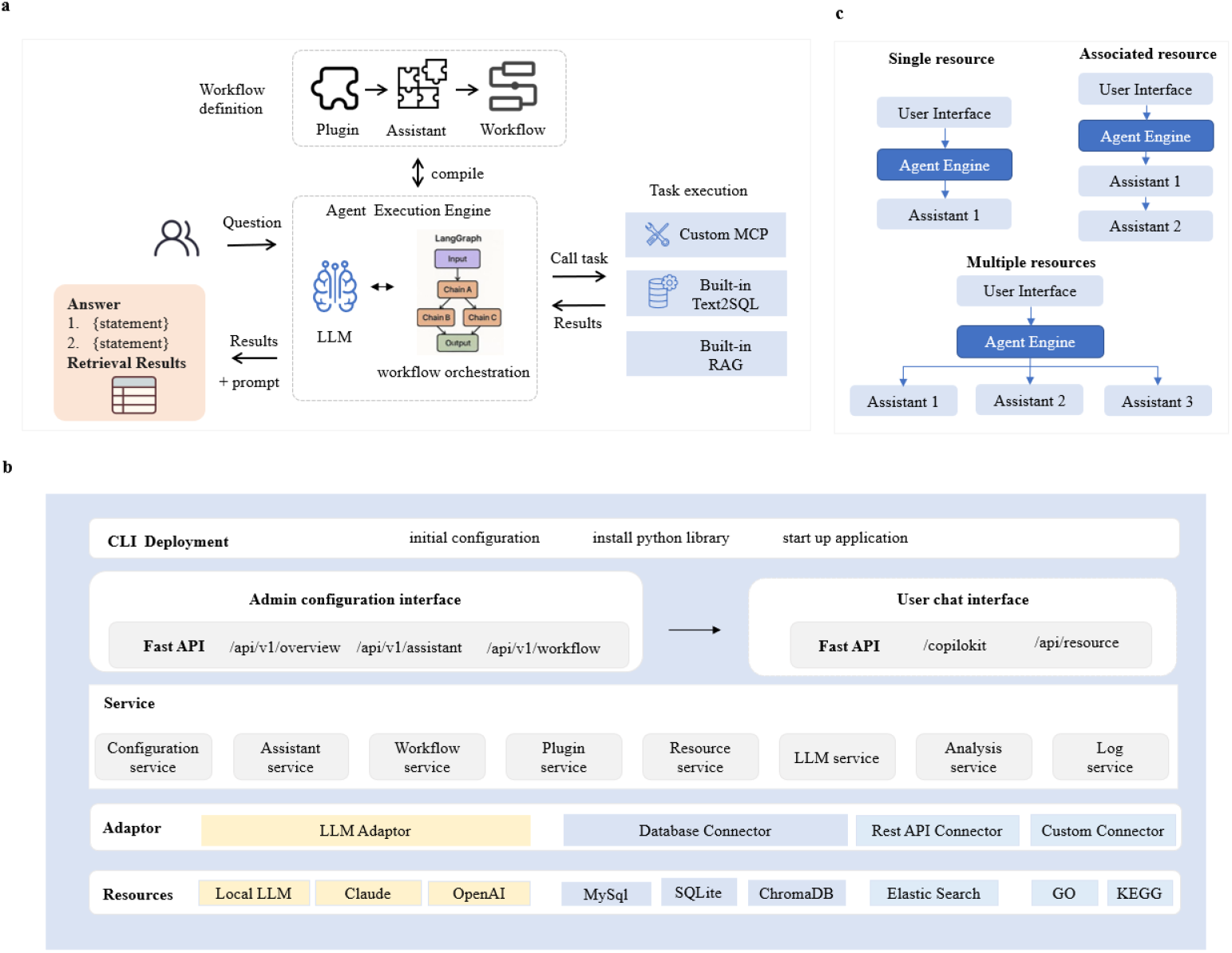
Overview of the study. a Overview of the Dingent framework. b Overview of the Dingent architecture. c Three application scenarios for data retrieval.

### The main features of Dingent

#### 1. One-stop agent application construction

Dingent modularizes the complex process of agent construction and provides an intuitive interface, enabling researchers without programming expertise to build customized agents through configuration. It offers a React-based visual management interface for configuring large language models, defining heterogeneous data source (such as MySQL, SQLite, ChromaDB, and Elastic Search), assembling plugins to construct assistants, and designing workflows to implement branching, conditional logic, and execution pathways. All configurations are stored in structured files. They are dynamically loaded at runtime and instantiated as executable agents. Through a react-based chat interface, users can interact with these agents by entering natural language queries to receive summarized answers along with traceable retrieval results. This integrated environment for configuration and interaction facilitates rapid, iterative improvement of agent applications.

#### 2. High-performance agent execution engine

Dingent is powered by a LangGraph-based ReAct execution engine, using asynchronous scheduling and caching to provide advanced natural language retrieval with high throughput and low latency. The engine’s design has three key aspects: First, upon receiving a user request, it dynamically constructs a workflow execution graph through an asynchronous graph-building mechanism. This avoids blocking other request and enables parallel execution of different nodes. Second, a double-checked caching structure is implemented, utilizing double-checked locking to minimize redundant workflow construction and initialization overhead. When the cache is hit, cached workflow can be rapidly reused. Finally, a reactive node execution strategy is implemented, in which each assistant functions as an independent node capable of dynamically selecting plugins, generating execution plans, and triggering data retrieval operations based on context. This design ensures both interpretability and adaptability throughout the execution process. In high-concurrency retrieval scenarios, this architecture effectively reduces redundant computations, accelerates response time, and provides a high-performance foundation for large-scale biological database retrieval tasks.

#### 3. Flexible building-block design for plugins, assistants, and workflows

The Dingent framework uses three core abstractions, namely plugins, assistants, and workflow, that together enable a flexible, building-block approach to constructing intelligent agent applications. Plugins serve as the minimal executable units, each designed to perform an atomic task, such as Text2SQL query generation, RAG retrieval, vector database queries, custom function calls (e.g. GO enrichment analysis), and remote service invocations (including MCP tools and REST APIs). Each plugin independently defines its input parameters, execution logic, and prompt templates, enabling the construction of reusable retrieval capabilities. These plugins can be further incorporated into assistants to form a complete functional unit. For instance, an assistant designed to answer questions about dogs’ phenotype and perform Gene Ontology (GO) enrichment analysis would simultaneously use Text2SQL, RAG, and a custom plugin for GO enrichment. The assistant defines its behavior within the workflow, including the order of execution, the structure of inputs and outputs, and strategies for error recovery. Furthermore, assistants can be assembled into more complex applications through workflows, a design based on a graph structure in which each assistant serves as a node. This architecture enables key functionalities including multi-assistant routing, conditional branching, node aggregation, and the orchestration of execution paths with dependency management. Completed workflows can be configurated and deployed for complex agent-based application such as multi-database federated retrieval and cross-database correlation analysis.

#### 4. Extensible plugin development

To accommodate diverse database types and user needs, Dingent features a flexible and extensive plugin architecture. It provides declarative dependency isolation, allowing users to specify required resources, such as dependency models, database connections, and external tools, directly within the configuration. These resources are automatically loaded during initialization with isolated environments to prevent dependency conflicts through fastmcp. A dedicated plugin lifecycle manager automates plugin discovery and registration, handling entry point scanning, metadata registration, and automatic attachment to the Assistant. It also supports execution hooks for pre- and post-processing, alongside mechanisms for rapid scheduling and error handling. Currently, the framework offers both built-in plugins, such as Text2SQL (for MySQL and SQLite) and RAG (fir ChromaDB and FAISS), and custom plugins based on the MCP protocol. With minimal configuration, users can seamlessly extend the system’s capabilities to build highly customized functions.

#### 5. Streamlined deployment, version management, and portability

Dingent is deployed as an installation package with a bundled front-end interface, allowing for one-click installation under Windows, Linux, and macOS. To ensure the reproducibility of built applications, Dingent provides version-locking and includes an automated migration mechanism that seamlessly updates existing configurations to newer versions. Once installed, two web service are launched: an administrative panel for configuration and a user chat interface. These features collectively enable rapid and portable deployment of Dingent across diverse research environments.

### Cases

#### Application 1: data retrieval in one single database through MCP

Dingent specialized in constructing dedicated agent application for a single database. This capability makes it ideal for the majority of common applications.

GenBase (10), a core database of NGDC (11), provides public access to nucleotide and protein data. To enable advanced search functionality, these datasets are separately indexed using Elasticsearch (ES), and queries are executed by invoking native ES functions via Java. Utilizing the SpringBoot MCP framework, we developed a MCP plugin for nucleotide and protein search. The plugin was then configured within the Dingent, where an assistant and a workflow were established. Following the launch of the user chat interface, this configuration allows users to automatically query nucleotide or protein sequences using natural language (Figure 2).

**Figure 2.**
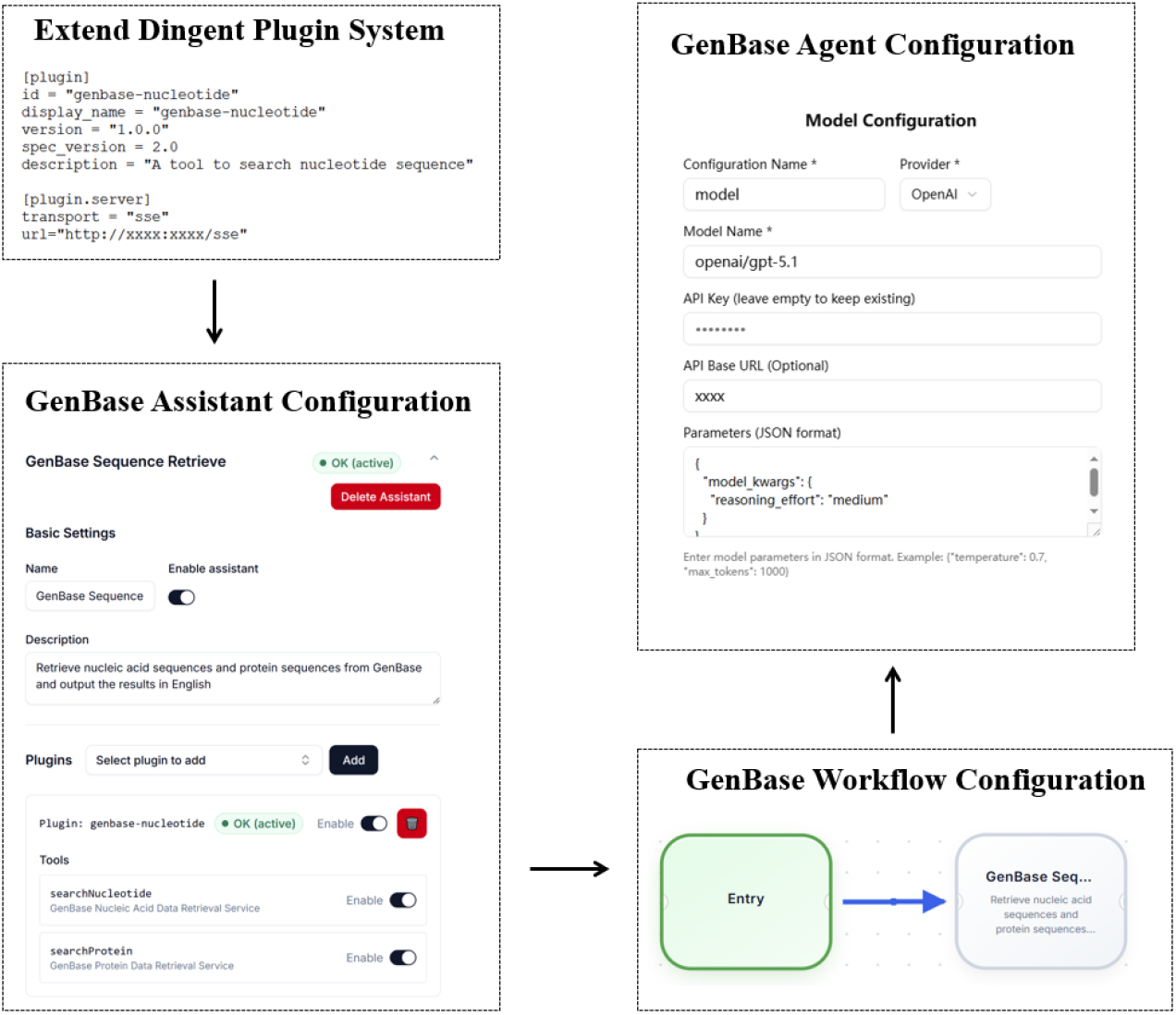
A flowchart of the GenBase search using the MCP tool via the Dingent framework

To retrieve the nucleotide sequences of *Ciona savignyi*, enter the query: “get nucleotide sequence of Ciona savignyi”. This will prompt Dingent to automatically invoke the nucleotide MCP service, returning and displaying the results on the webpage. The output indicates 88,120 available nucleotide sequences for *Ciona savignyi*, and the first 24 sequences show as a table in the middle panel, a summarized results from LLM show in the right panel. Figure 2 illustrates the GenBase search flowchart using the MCP tool in Dingent, and Figure 3 displays the corresponding results.

**Figure 3.**
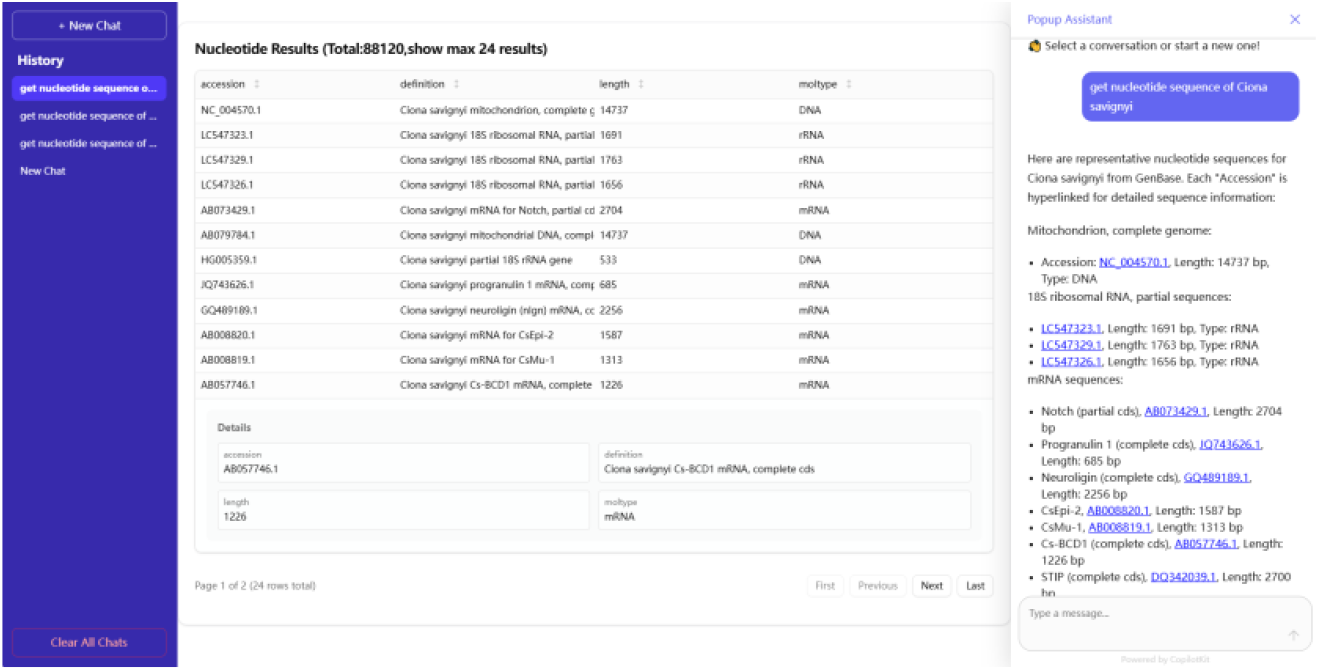
A screenshot of the *Ciona savignyi* Q&A result

#### Application 2: intelligent routine among multiple databases

Dingent enables the integration of multiple resources and intelligently routes user queries across them. This is particularly useful for users who need to integrate multiple resources into a unified agent platform.

Three resources including BioKA (12), GenBase and iDog (13) are configured as assistants in the admin panel. BioKA and iDog leverage MySQL for data storage, employing the built-in Text2SQL and RAG plugins, whereas GenBase uses a custom plugin. Using the workflow configuration panel, a branch selection option is established (Figure 4). Once the front-end interface is launched, queries like “Which dog breeds require daily grooming?” are automatically sent to iDog, while “Search for the biomarker TP53.” are routed to BioKA. This result underscores the reliability of the proposed framework.

**Figure 4.**
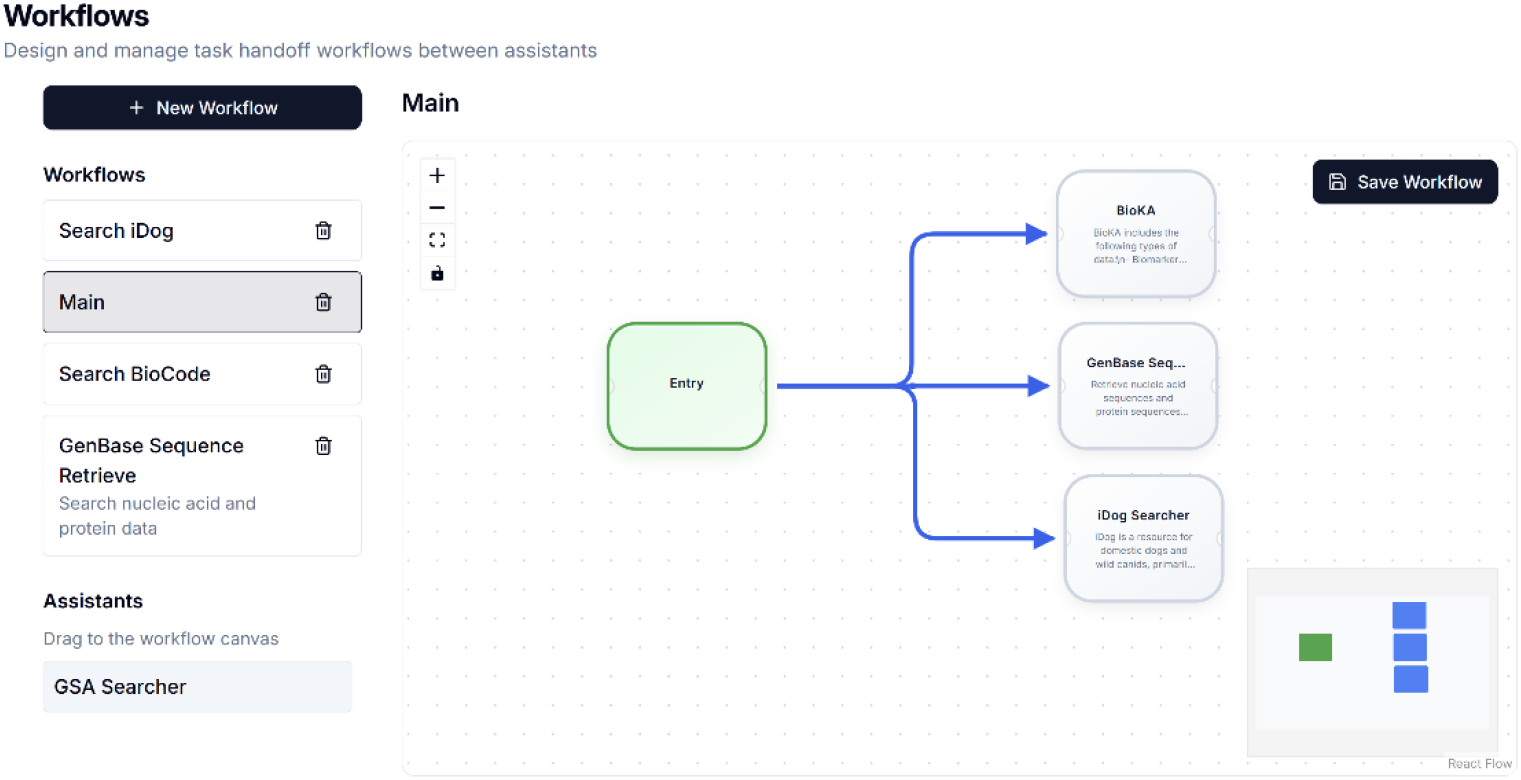
A workflow chart for multiple resources configuration

**Table 1.** Partial curated questions.

| Questions | Data resource |
| --- | --- |
| Search for the biomarker TP53. | BioKA |
| Retrieve information on the IGF2 biomarker. | BioKA |
| Look up the JAK2 biomarker. | BioKA |
| Search for biomarker data on MSH5. | BioKA |
| Find biomarkers associated with diabetes. | BioKA |
| List dog breeds prone to hereditary heart disease. | iDog |
| Which dog breeds require daily grooming? | iDog |
| Are Poodles prone to epilepsy, and is it associated with the SCN1A gene? | iDog |
| What genetic factors are linked to Gastric Dilatation-Volvulus (GDV) in large dog breeds? | iDog |
| Are Labrador Retrievers susceptible to Hip Dysplasia, and what are the key genes involved? | iDog |
| Retrieve studies with titles containing "whole genome sequencing" | GenBase |
| Retrieve genetic markers related to diabetes | GenBase |
| Retrieve transcription factors containing "zinc finger" domains | GenBase |
| Retrieve records with author affiliation containing "Harvard University" | GenBase |
| Retrieve gene clusters related to antibiotic biosynthesis | GenBase |

#### Application 3: associated query of dog disease-associated gene and reported biomarker

Dingent can handle queries across two or more data resources, making it particularly useful when the results are interrelated.

iDog houses disease and gene associations, and BioKA contains gene biomarkers. Within the Dingent framework, a connection between iDog and BioKA is established in the workflow panel, enabling AI-driven applications for the identification of gene markers for dog disease (Figure 5).

**Figure 5.**
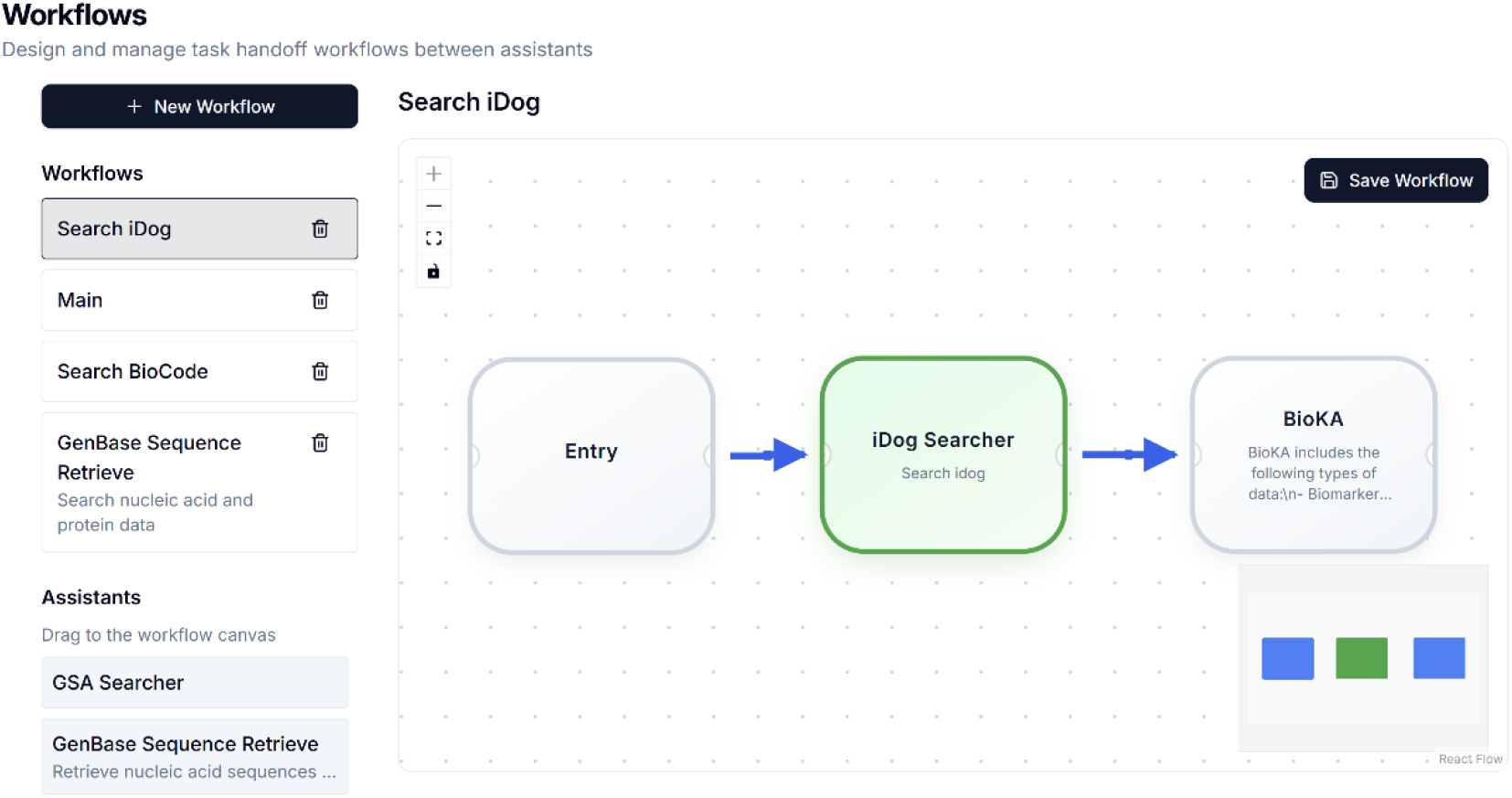
A workflow chart for associated resources configuration

Canine degenerative myelopathy (DM) has been associated with mutations in the superoxide dismutase 1 gene (SOD1) (14, 15), targeted SOD1 gene silencing (e.g., via siRNA) has identified several functional leads that could be evaluated in a veterinary clinical trial in canine DM models (16). We aim to identify genes associated with DM and investigate their potential as biomarkers. When processing a query such as “Give me disease cause information and causal genes of degenerative myelopathy in domestic dogs”, Dingent searches the iDog and identifies two causal genes SOD1 and SP110. If a follow-up query asks, “Tell me which are biomarkers for this disease in the previous causal genes”, Dingent uses SOD1 and SP110 to search the BioKA. After filtering the results, it returns the answer “Biomarker for this disease among the previous causal genes: SOD1 (protein; diagnostic biomarker)”. This finding is consistent with the previous reports. A screenshot of the results is presented in Figure 6.

**Figure 6.**
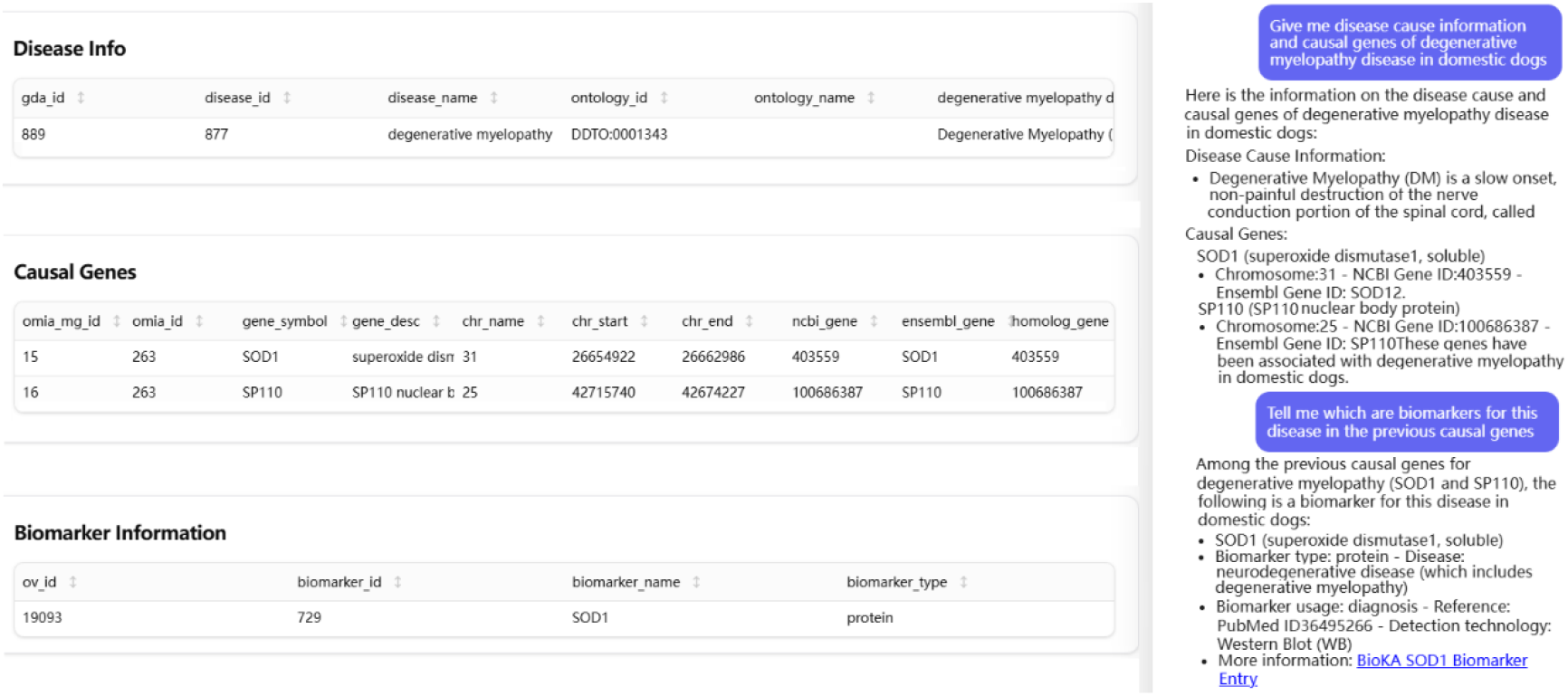
A screenshot of casual gene in degenerative myelopathy for iDog and BioKA

## Discussion and Conclusion

Dingent serves as a configurable, one-stop agent framework to provide an LLM-powered agent for various databases and a front-end web interface for natural language-based data retrieval. Its modular architecture, straightforward configuration process, and diverse data source access methods enhance data integration capabilities while offering the flexibility to extend additional function modules. Distributed as an open-source framework with user-friendly installation and configuration guides, Dingent is designed to be both extensible and easy to deploy, making it a powerful tool for transforming custom data resource into AI-augmented resources through LLM-agent integration. To demonstrate its capabilities, we developed three applications using the framework: nucleotide sequence retrieval, intelligent routine among multiple databases, and querying disease-associated genes and biomarkers in dogs. Although the current applications are in life sciences, Dingent can be readily adapted to other fields, such as earth sciences and ecology, for efficient data discovery.

Although Dingent incorporates a number of advanced capabilities, it is subject to several important limitations. First, while Dingent employs individual assistants as autonomous agents, the framework currently lacks robust mechanisms for inter-agent collaboration and communication, which would be essential for complex, multi-step tasks. Second, the system does not include a user management module, limiting its applicability in scenarios that require secure user authentication and role-based access control. Third, the current functionality of Dingent is predominantly focused on data retrieval, whereas its support for data analysis is less developed.

In the future, we will continuously enhance Dingent by integrating a wider variety of plugins and introducing a plugin management panel to simplify discovery and installation. For the front-end web interface, we will add features like user session management and multi-language support. Furthermore, we plan to integrate a deep search function powered by AI, which will provide more flexible data search and analysis strategies through intelligent planning.

## Code availability

The codes developed in this study are openly accessible on GitHub at https://github.com/saya-ashen/Dingent. The code has been packaged, allowing for easy installation by following the instructions provided at home web page https://ngdc.cncb.ac.cn/dingent.

